# Mitochondrial dysfunction compromises ciliary homeostasis in astrocytes

**DOI:** 10.1101/2021.08.24.457472

**Authors:** Olesia Ignatenko, Satu Malinen, Helena Vihinen, Joni Nikkanen, Aleksandr Kononov, Eija Jokitalo, Gulayse Ince-Dunn, Anu Suomalainen

**Affiliations:** Stem Cells and Metabolism Research Program, Faculty of Medicine, University of Helsinki, 00290 Helsinki, Finland; Institute of Biotechnology, University of Helsinki, 00014 Helsinki, Finland; Cardiovascular Research Institute, University of California, 94143-0795 San Francisco, USA; Cancer Research UK, University of Manchester, SK10 4TG Manchester, UK; Neuroscience Center, University of Helsinki, 00290 Helsinki, Finland; HUSlab, Helsinki University Hospital, 00290 Helsinki, Finland

## Abstract

Astrocytes, often considered as secondary responders to neurodegenerative processes, are emerging as primary drivers of brain disease. The underlying pathogenic mechanisms are, however, insufficiently understood. Here we show that pathogenesis of mitochondrial spongiotic encephalopathy, a severe manifestation of mitochondrial brain diseases, involves abnormal maintenance of the astrocytic primary cilium, a major signaling organelle of a cell. We show that progressive respiratory chain deficiency in astrocytes activates FOXJ1 and RFX transcription factors and master regulators of motile ciliogenesis. Consequently, a wide aberrant nuclear expression program with FOXJ1 and RFX target genes, encoding motile cilia components, is induced in astrocytes. While the affected astrocytes still retain a single cilium, these organelles elongate and become remarkably distorted. Multiciliated ventricle-lining ependymal cells show no overt cilia morphology defects despite similar mitochondrial dysfunction. We propose that the chronic activation of the integrated mitochondrial stress response (ISR^mt^), specifically induced in astrocytes, drives anabolic metabolism and promotes ciliary growth. Collectively, our evidence indicate that 1) an active signaling axis exists between astrocyte mitochondria and primary cilia; 2) ciliary signaling is part of ISR^mt^ in astrocytes; 3) metabolic ciliopathy is a novel pathomechanism for mitochondria-related neurodegenerative diseases.

## Introduction

Astrocytes are essential for the homeostasis of the central nervous system (CNS), maintaining ionic balance, blood-brain barrier integrity, synapse function, and the metabolic homeostasis (Sofroniew and Vinters, 2010; Wang and Bordey, 2008). When stressed by disease, injury or metabolic insults, astrocytes transition into a state termed reactive astrogliosis, characterized by considerable functional, morphological, and molecular remodeling. For example, reactive astrocytes contribute to inflammatory and vascular responses, scar tissue formation, and metabolic support of the CNS cells. Depending on the insult, these responses may be protective or toxic (Anderson et al., 2016; Bush et al., 1999; Liddelow et al., 2017; Yun et al., 2018). Despite the essential functions of astrocytes in CNS metabolism, the molecular programs induced by metabolic distress remain insufficiently understood.

Mitochondrial dysfunction is a hallmark of neurodegeneration, characteristic for primary mitochondrial diseases (Gorman et al., 2016) and also secondarily implicated in common neurological diseases, such as Parkinson’s and Alzheimer’s diseases, amyotrophic lateral sclerosis, and Huntington’s disease (Lin and Beal, 2006; Wang et al., 2019). Primary mitochondrial diseases manifest, however, with a large variation of different disease entities, indicating that mitochondrial mechanisms underlying neurological disease manifestations depend on the type of dysfunction and may be cell-type-specific. Elucidation of pathophysiological mechanisms in primary mitochondrial models is important to understand the variable roles of mitochondria for neurodegeneration in general.

Recent data highlight astrocytes as drivers of pathogenesis of mitochondrial spongiotic brain diseases, typical for children’s devastating and incurable encephalopathies (Ignatenko et al., 2018). Intriguingly, mitochondrial dysfunction specifically in astrocytes induces reactive astrogliosis, which was sufficient to drive severe brain pathology in mice (Ignatenko et al., 2018, 2020; Murru et al., 2019). Loss of mitochondrial DNA helicase Twinkle (Twnk) in astrocytes (TwKO^astro^; KO for knockout) and consequent progressive depletion of mitochondrial DNA (mtDNA) leads to pervasive reactive astrogliosis starting after two months of age, with progressive vacuolation of brain parenchyma, consequent reactive microgliosis, myelin disorganisation, and premature death (Ignatenko et al., 2018). The brain pathology of TwKO^astro^ mice closely resembles human spongiotic encephalopathies caused by mtDNA maintenance defects (Kollberg et al., 2006; Palin et al., 2012). The brain of TwKO^astro^ mice undergoes metabolic reprogramming, including the induction of a mitochondrial integrated stress response (ISR^mt^) (Ignatenko et al., 2020). Twnk deletion and mtDNA loss in cortical neurons is tolerated until seven months of age, after which the mice manifest with an acute-onset neurodegenerative disease (Ignatenko et al., 2018), but show no signs of ISR^mt^ (Ignatenko et al., 2020).The astrocytic ISR^mt^ components resemble, but not completely overlap with responses in other models with mitochondrial insults (Bao et al., 2016; Forsström et al., 2019; Khan et al., 2017; Kühl et al., 2017; Mick et al., 2020; Murru et al., 2019; Nikkanen et al., 2016; Richter et al., 2013; Tyynismaa et al., 2010). The tissue- and cell-type specific ISR^mt^ components are excellent candidates to explain the highly variable tissue-specific manifestations of mitochondrial diseases. Therefore, understanding physiological consequences of metabolic stress responses in astrocytes is essential, as these cells have major metabolic functions in the nervous system.

Here we report that astrocytes with mitochondrial respiratory dysfunction show anomalous induction of a motile ciliogenesis program and major lipid metabolic remodeling. The ciliary response is mediated by transcription factors RFX (Regulatory Factor X) family and FOXJ1 (Forkhead Box J1), with no previously known roles in adult astrocytes or ISR^mt^. Astrocytes possess primary cilia (Kasahara et al., 2014; Sipos et al., 2018; Sterpka and Chen, 2018), but little is known about their roles in disease. Our findings of a major ciliogenic response to mitochondrial respiratory dysfunction in astrocytes constitute an exciting new front for astrocyte research in the context of brain pathologies, with special relevance for mitochondrial spongiotic encephalopathies of children.

## Results and discussion

### Astrocytes lacking Twinkle helicase develop mtDNA depletion

To investigate the astrocyte-specific metabolic responses to mitochondrial dysfunction, we purified cortical astrocytes from 3-3.5 months old TwKO^astro^ and control mice, using magnetic beads coated with antibodies against astrocyte-specific surface antigen ATP1B1 (ACSA-2) (Batiuk et al., 2017). At this age, TwKO^astro^ mice show early-stage disease, with mild gliosis and sparse vacuoles in brain parenchyma (Fig. S1), but advanced pathological changes, such as myelin disorganization or neuronal loss (Ignatenko et al., 2018), are not yet observed. Astrocytes were enriched in the ACSA-2^+^ fraction (RT-qPCR analysis of cell-type-specific markers; Fig. 1A). The astrocytes from TwKO^astro^ mice showed a profound mtDNA depletion, which was not detectable in total brain lysates (Fig. 1B). This is a consequence of successful inactivation of Twinkle, the replicative mtDNA helicase.

**Fig.1:**
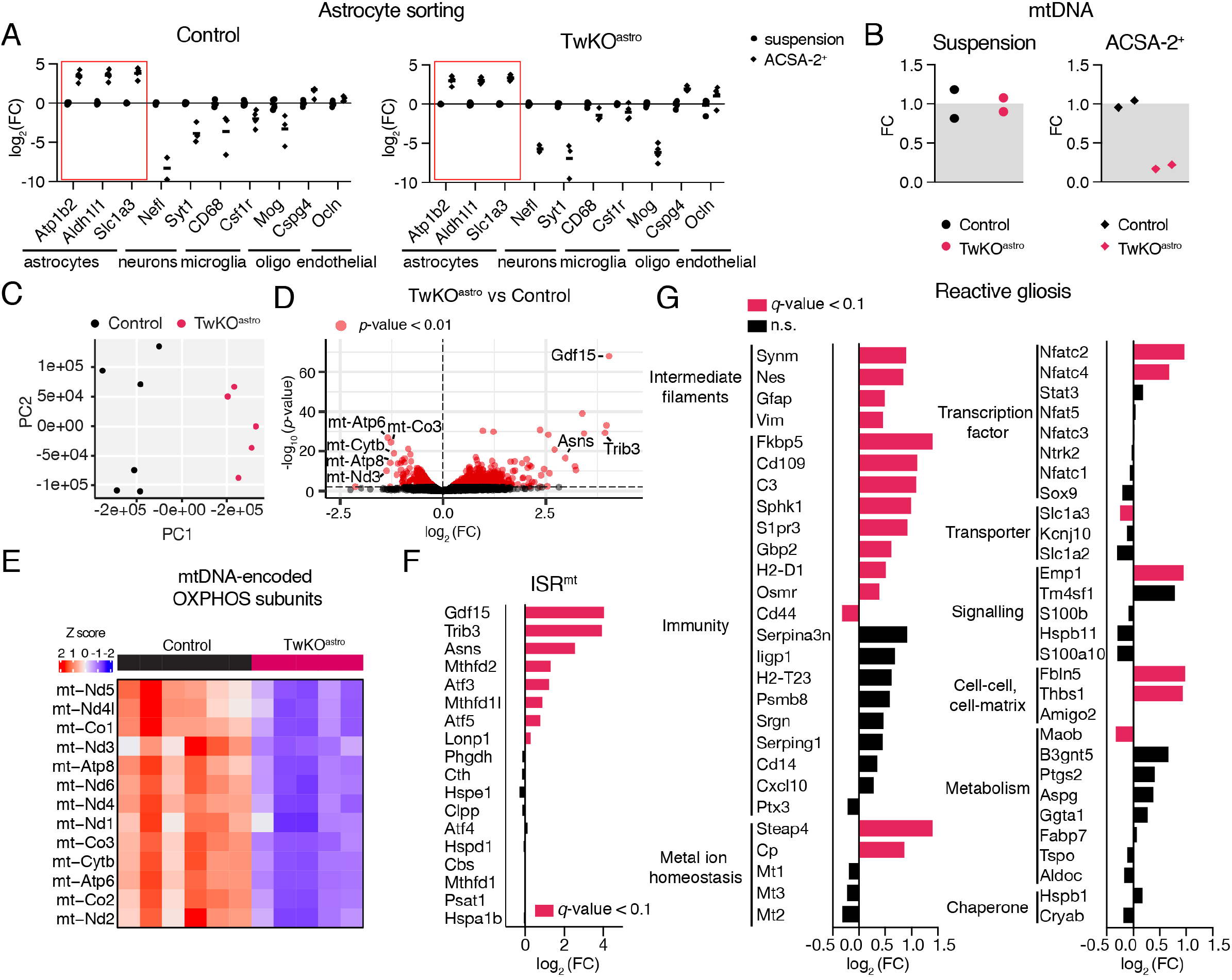
mtDNA depletion in astrocytes leads to cellular stress responses. (A) Enrichment of cell-specific genes in ACSA-2^+^ fraction compared to unsorted suspension, measured using RT-qPCR. Genes of interest are normalised to Hmbs. (B) MtDNA amount, normalised to nuclear DNA. Measured using qPCR. (C) Transcriptome of astrocytes, principal component (PC) analysis. (D) Transcriptome of astrocytes, all identified genes. Gene symbols in the volcano plot denote those most downregulated genes that are encoded in the mtDNA and those most upregulated genes that are ISR^mt^ components (red dots indicate p-value < 0.01). (E) Heatmap of mtDNA transcripts. (F) Transcripts encoding ISR^mt^ components (gene list is curated based on (Forsström et al., 2019). (G) Transcripts that mark reactive astrogliosis (gene list is curated based on (Escartin et al., 2021; Liddelow et al., 2017; Zamanian et al., 2012)). See also Suppl. Table 1. (C-H): RNA sequencing, astrocytes purified from Ctrl (n=6) and TwKO^astro^ (n=5) mouse brain cortical preparations, 3-3.5 months old mice. See also Suppl. Table 6. FC = fold change; ISR^mt^ = mitochondrial integrated stress response; ACSA-2 = astrocyte cell surface antigen-2; qPCR = quantitative polymerase chain reaction; RT-qPCR = real-time qPCR.

Unsupervised principal component analysis of RNAseq data, derived from the isolated astrocytes, showed that control and TwKO^astro^ samples clustered separately (Fig. 1C). Expression levels of 1,131 genes were upregulated and 408 were downregulated (|log_2_(FC)|>0.3 and q-value<0.1) (Fig. 1D). Consistent with mtDNA depletion, mtDNA-encoded transcripts were among the most down-regulated (Fig. 1D, E). These results provided a proof of principle that the purified fraction represented the astrocyte population with Twnk-KO.

### Astrocytes lacking Twinkle induce a partial ISR^mt^ and molecular markers of reactive astrogliosis

A close examination of cellular stress response genes revealed that several defining components of ISR^mt^ were induced in astrocytes purified from TwKO^astro^: activating transcription factors (Atf3, Atf5) and Trib3 kinase that govern the response; mitochondrial folate metabolism genes (Mthfd2, Mthfd1l); Asns that catalyses conversion of aspartate to asparagine, and the metabolic hormone Gdf15 (Fig. 1D, F). Previously, the mRNAs of the rate-limiting enzymes of serine biosynthesis (Psat1, Phgdh) and transsulfuration (Cth, Cbs) were reported to be increased in the cortical lysates of 5 months old TwKO^astro^ mice (Ignatenko et al., 2020). These two pathways were not yet induced at 3-3.5 months timepoint in our purified astrocytes or cortical lysates of TwKO^astro^ mice (Fig. 1D). The stage-wise induction of ISR^mt^ in astrocytes mimics that in the skeletal muscle, in a mouse model expressing a dominant Twinkle mutation (Forström et al., 2019). However, the components of ISR^mt^ are partially different in the astrocytes and the muscle. For example, FGF21, a major regulator and endocrine signaler of ISR^mt^ in the muscle, is not part of the astrocyte response. These results highlight the cell-type-specificity of these responses.

We then examined markers of reactive astrogliosis previously published to be changed in various brain pathologies (Suppl. Table 1) (Escartin et al., 2021; Liddelow et al., 2017; Zamanian et al., 2012). Out of 56 such marker genes, expression levels of 22 were changed in TwKO^astro^ (Fig. 1G). Genes encoding intermediate filaments and factors involved in immune responses (e.g. nuclear factor of activated T cells; Nfatc2 and Nfatc4), were most robustly upregulated in our dataset (Fig. 1G). These data show that mitochondrial pathology causes astrocyte reactivity that shares components with responses caused by other nervous system stresses.

### Mitochondrial dysfunction in astrocytes induces a unique ciliogenic program

Intriguingly, the top five upregulated pathways in TwKO^astro^ astrocytes were related to ciliary functions (Fig. 2A). The majority of differentiated eukaryotic cells, including astrocytes, possess a primary signalling cilium (Dahl, 1963; Karlsson, 1966; Kasahara et al., 2014; Sterpka and Chen, 2018). Surprisingly, the ciliary pathways upregulated in TwKO^astro^ astrocytes were related to motile cilia (Fig. 2A) that facilitate liquid movement at the surface of lumen-facing cells, while a primary cilium is specialized to sense and integrate external signals critical for cell proliferation and differentiation. To explore the extent of the pathway induction in TwKO^astro^ dataset, we used SysCilia and CiliaCarta databases (van Dam et al., 2013, 2019). TwKO^astro^ astrocytes showed a remarkable upregulation of 64 out of 280 SysCilia and 163 out of 791 CiliaCarta genes, while only four and 13 genes, respectively, were downregulated (Fig. 2B, Suppl. Table 2). These results indicated that mitochondrial dysfunction induces a striking and robust activation of motile ciliary program in astrocytes that harbor single immotile cilia.

**Fig.2:**
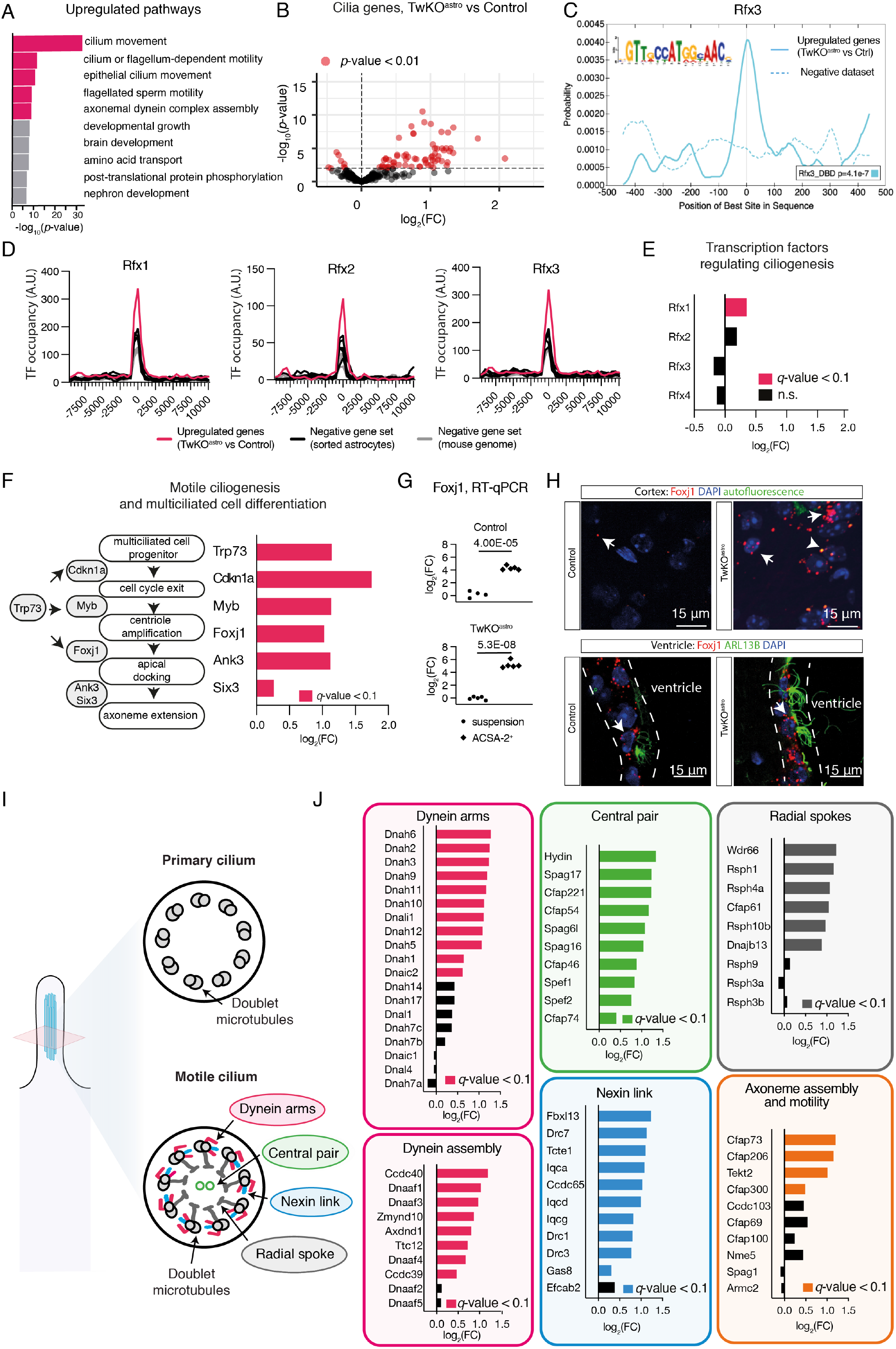
Mitochondrial dysfunction in astrocytes sorted from TwKO^astro^ mice induces motile ciliogenesis program. (A-F, J): RNA sequencing, astrocytes purified from TwKO^astro^ compared to control mice. The dataset is described in Fig. 1. (A) Gene ontology pathway enrichment analysis of upregulated genes in astrocytes of TwKO^astro^ mice. Pathways involved in cilia motility are marked in red. (B) Genes expressing ciliary proteins in astrocytes of TwKO^astro^ mice (SysCilia Gold Standard gene list (van Dam et al., 2019)). See also Fig. S2A and Suppl. Table 2. (C) Motif enrichment analysis of promoters of upregulated genes in TwKO^astro^ astrocytes reveal RFX consensus motif (RFX3 motif enrichment is shown here, see also Fig. S2B). (D) Distribution graphs of the TF binding sites proximal to the promoters of upregulated genes (ChIP-seq data from (Lemeille et al., 2020)). X-axis represents distance from TSS (transcription starting site), which equals point zero. Randomly selected gene sets either from the whole mouse genome or genes expressed in astrocytes, but not changed in our dataset, constitute negative gene sets. See also Suppl. Table 3. (E) Transcription factors that regulate ciliogenesis in the brain; mRNA levels in TwKO^astro^ astrocytes. (F) Key steps of multiciliated cell differentiation, from cell cycle exit to motile cilia formation; schematic representation (left). The expression of the key regulators of cilia biogenesis in astrocytes of TwKO^astro^ mice. (G) Foxj1 expression in purified astrocytes (ACSA-2^+^ fraction) compared to unsorted cell suspension; RT-qPCR, normalised to Ywhaz transcript level. Symbols represent individual preparations. p-values calculated using unpaired two-tailed parametric t-test. (H) Foxj1 expression; RNA-fluorescence in situ hybridization, 3.5 months old mice. Dashed lines indicate the ependymal cell layer. Puncta specific for Foxj1 channel were analysed (arrows). Puncta that overlap with autofluorescence or ARL13B signal (arrowhead) were excluded as nonspecific signal. Mice also express AAV-gfaABC1D-Arl13b-eGFP to visualize cilia. See also Fig. S2D-E. (I) Schematic representation of cilia cross-section. Axonemes of both primary and motile cilia comprise nine doublet microtubules. Motile cilia also harbor components that are not present in primary cilia. (J) Expression of structural components specific to motile cilia and other factors involved in axoneme assembly and motility in astrocytes of TwKO^astro^ mice compared to control mice.

### Mitochondrial dysfunction in astrocytes induces expression of motile cilia components via RFX and FOXJ1 transcription factors

Next, we asked whether specific transcription factor (TF) consensus sequences were enriched in the promoters of genes upregulated in astrocytes from TwKO^astro^ mice (1,000 bp window centering on the transcription start sites). The analysis revealed an enrichment of motifs recognized by RFX TF family (Fig. 2C, S2B). Intriguingly, RFX1-4 TFs have been described as master regulators of ciliogenesis in a wide range of cell types (Santos and Reiter, 2008; Thomas et al., 2010). To investigate whether experimentally determined RFX binding sites were also enriched in TwKO^astro^ dataset, we used publicly available RFX1-3 ChIP-Seq datasets derived from mouse multiciliated ependymal cells (Lemeille et al., 2020). This analysis revealed that RFX binding sites were significantly over-represented in genomic regions proximal to transcription start sites of genes upregulated in TwKO^astro^ astrocytes compared to negative gene sets (Fig. 2D, Suppl. Table 3). Rfx1 transcript level was slightly increased (log_2_(FC)=0.35, q-value=3.0E-03), while other RFX family member mRNAs were unchanged in TwKO^astro^ dataset (Fig. 2E, S2C). These findings suggest that in addition to a slight RFX1 induction, RFX-family transactivation or their recruitment to promoters might be activated in our astrocyte population. Furthermore, our systematic search for ciliogenesis regulators led to discovery of an upregulation of Trp73 and Foxj1, two TFs that control motile ciliogenesis and differentiation of multiciliated cells (Trp73 log_2_(FC)=1.15, q-value=5.56E-08; Foxj1 log_2_(FC)=1.5, q-value=1.02E-06) (Fig. 2F, S2C). Trp73 encodes TP73, which triggers a downstream cascade towards multiciliated cell differentiation, including FOXJ1 (Napoli and Flores, 2016; Nemajerova et al., 2016). We found key downstream TP73 targets (Cdkn1a, Myb, Foxj1, Ank3 and Six3) to be all upregulated in our dataset (Fig. 2F). In summary, the transcriptional response to mitochondrial dysfunction in astrocytes converged on a robustly activated motile ciliogenic program.

FOXJ1 is a master regulator of motile ciliogenesis that works in coordination with RFX TFs in the brain and is presumed to be active only in ependymal cells (Jacquet et al., 2009; Stubbs et al., 2008; Yu et al., 2008), the only multiciliated cell type in the central nervous system. However, Foxj1 expression in sorted astrocytes was markedly higher than in unsorted brain suspension both in control and TwKO^astro^ mice, suggesting that Foxj1 is expressed in adult astrocytes and upregulated as a consequence of mitochondrial stress (Fig. 2F, G). Next, we asked whether Foxj1 was indeed upregulated in cortical astrocytes or only in ependymal cells, which could co-purify with astrocytes in our isolation protocol. The spatial analysis of Foxj1 expression (RNA-fluorescence in situ hybridization) in TwKO^astro^ cerebral cortex showed markedly increased expression at 3.5 months of age in comparison to controls (Fig. 2H, S2D). Foxj1-positive puncta were present also in the cortex of control mice, further suggesting that the expression of Foxj1 is not restricted to ependymal cells in the brain (Fig. 2H, S2E). As expected, ependymal cells, but not subventricular zone cells, showed a strong Foxj1 signal (Fig. 2H). Our evidence suggests that FOXJ1, a regulator of multiciliated cell differentiation, is not only part of a developmental program of ependymal cell differentiation, but is also expressed in the cerebral cortex of adult mice, and its expression is upregulated in astrocytes as a response to mitochondrial dysfunction.

To investigate the extent of activation of a motile cilia program in TwKO^astro^, we curated a catalogue of genes that encode proteins that are either unique to motile cilia, unique to primary cilia or are pan-ciliary (Suppl. Table 2). Axonemes of both primary and motile cilia comprise nine outer doublet microtubules, while motile cilia also harbor a set of distinct components essential for their movement (Fig. 2I). Expression of genes that compose all key structures specific to motile cilia were induced in TwKO^astro^ astrocytes (Fig. 2J). These included dynein arms and the cytoplasmic factors required for their assembly; the nexin-dynein regulatory complex (nexin link) that is located between microtubule doublets; the central pair (an additional microtubule doublet); the radial spokes that protrude towards the central pair; and other factors involved in the assembly and motility of the axoneme (Fig. 2J). The expression of most genes not specific to motile cilia (primary cilia or pan-ciliary genes) were unchanged (Fig. S2F).

Ectopic expression of FOXJ1 in Xenopus and Danio rerio is sufficient to induce formation of functionally motile cilia in cells normally devoid of these organelles (Stubbs et al., 2008; Yu et al., 2008). Our data show that mitochondrial respiratory dysfunction in the adult mouse brain upregulates Foxj1 and the motile ciliogenic program in astrocytes, a cell type that normally possesses a single primary cilium.

### Motile ciliogenesis program is specific to metabolic and growth factor-related astrocyte stress

The anomalous motile ciliogenesis and multiciliated cell differentiation program induction in TwKO^astro^ astrocytes prompted us to explore whether such response is typical also for other astrocyte stress models (Anderson et al., 2016; Guttenplan et al., 2020; Li et al., 2019), ageing astrocytes (Boisvert et al., 2018), or other mouse models with mitochondrial dysfunction (Kühl et al., 2017). While in TwKO^astro^ astrocytes, 62 out of 92 genes encoding motile cilia components or ciliogenic or multiciliated cell differentiation factors were changed compared to controls (Figure S2G, Suppl. Table 2), other mtDNA maintenance KO models did not show such changes. For example, ciliogenic program was not induced in the cardiac muscle of heartspecific Twnk KO mice (Kühl et al., 2017), signifying astrocyte specificity of the response (Figure S2G). Cultured astrocytes inhibited for epidermal growth factor signalling (EGFR silencing or HBEGF withdrawal) displayed a robust upregulation of the motile ciliogenesis program, including induction of Foxj1 (Figure S2G) (Li et al., 2019). In contrast, reactive astrocytes stimulated with pro-inflammatory IL1+TNF**α**+C1q cocktail or sorted from mice with spinal cord injury displayed partial downregulation of the motile ciliogenesis program (Figure S2G) (Guttenplan et al., 2020). Finally, astrocytes sorted from aged mouse brain showed no changes (Figure S2G) (Boisvert et al., 2018). These data suggest that mitochondria – cilia communication axis is cell-type specific and responds to mitochondrial dysfunction.

### Astrocytic cilia are elongated and contorted in TwKO^astro^ mice

The robust upregulation of the motile multiciliary program in Twnk knockout astrocytes prompted us to ask whether morphology of primary and/or motile cilia is altered in TwKO^astro^ mice. Even in mice with advanced disease (4.5-5 months; spongiotic pathology, wide-spread reactive astrogliosis; Fig. S1) (Ignatenko et al., 2018) the astrocytes were monociliated, as determined by immunofluorescent staining with the ciliary axoneme protein ARL13B together with GFAP or ALDH1L1, two markers for reactive astrocytes (Fig. 3A, 3B, respectively). Cells positive for either marker revealed a shift in length distribution towards long cilia in TwKO^astro^ mice: 2.0 - 6.5 μm in controls and 2.5 - 8.2 μm in TwKO^astro^ cortex (Fig. 3C). To better analyse cilia shapes, we classified them as straight, bent or contorted (Fig. 3D). Contorted cilia in TwKO^astro^ included S-shaped and corkscrew-like morphologies, and occasionally the long cilia appeared to form several loops (Fig. 3F, bottom panel). Such extreme morphologies never occurred in control mice. Generally, the proportion of cilia with contorted morphology was higher in TwKO^astro^ than in controls (Fig. 3E). Finally, among factors contributing to primary cilia signaling, the expression of the sonic hedgehog pathway effector Gli1 was induced in astrocytes sorted from TwKO^astro^ mice (Fig. S2F). These results indicate that mitochondrial dysfunction in astrocytes remarkably modifies the structure of the primary signalling organelle of the cells, the cilium.

**Fig.3:**
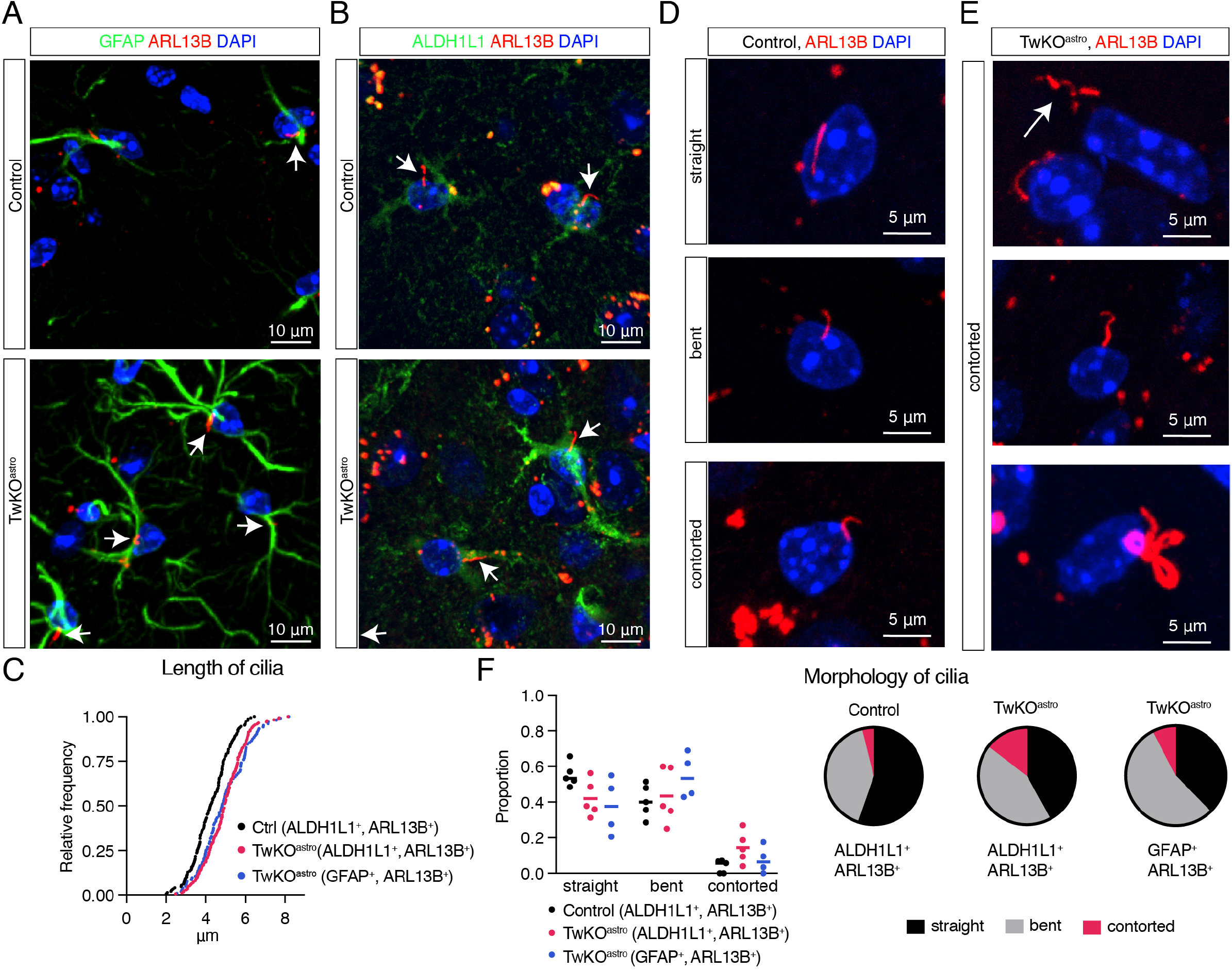
Astrocyte cilia elongate and become contorted upon astrocytic mitochondrial dysfunction. (A-F): Cerebral cortex, 4.5-5 months old mice. (A-B) Ciliary marker ARL13B immunostaining in GFAP^+^ (A) or ALDH1L1 ^+^ (B) astrocytes in control and TwKO^astro^ mice. (C) Length distribution of cilia in TwKO^astro^ and control mice. ARL13B signal in ALDH1L1^+^ or GFAP^+^ cells analysed. n=5 mice per genotype, 6 fields of view per mouse. Total number of cilia: control ALDH1L1^+^, ARL13B^+^ = 148; TwKO^astro^ ALDH1L1^+^, ARL13B^+^ = 174; TwKO^astro^ GFAP^+^, ARL13B^+^ = 75; control GFAP^+^, ARL13B^+^ cells were too few for analysis. Control vs TwKO^astro^ (ALDH1L1^+^ARL13B^+^): Kolmogorov-Smirnov test p-value < 0.0001, D = 0.5703; oneway analysis of variance: Pr (>F) = 6.76e-09. Dots represent individual cilia. (D) Representative images of cilia morphology in control mice. Immunostaining against ARL13B. (E) Cilia are contorted and elongated in TwKO^astro^ mice, immunostaining against ARL13B. (F) Quantification of cilia morphology in TwKO^astro^ and control astrocytes, based on the dataset from (C). Dots represent an average per mouse. Pie charts show average per genotype.

### Ependymal multiciliated cells with mitochondrial dysfunction show normal cilia structure

Next, we asked whether the multiciliary program in TwKO^astro^ mice also affected the structure of multiciliated ependymal cells. Initially, using the Rosa26-CAG-LSL-tdTomato reporter mice, we determined that GFAP-73.12-Cre driver line used to generate TwKO^astro^ mice was active in ependymal cells (Fig. S3A). These cells in 5-month-old TwKO^astro^ mice showed a marked deficiency of the activity of cytochrome c oxidase (the respiratory chain complex IV, partially encoded by mtDNA), whereas the activity of fully nuclear-encoded complex II was preserved, similarly to the findings in cortical astrocytes of TwKO^astro^ mice (Fig. S3B) (Ignatenko et al., 2018). Anti-ARL13B immunostaining indicated that the general morphology of ciliary tufts was preserved in TwKO^astro^ mice (Fig. 4A). Scanning electron microscopy also revealed normal ciliation across the ventricular wall (Fig. 3B). These results indicate that the aberrant ciliary structure was specific to primary cilia in astrocytes and that TwKO^astro^ had no apparent effects to motile cilia of the ependymal cells of the lateral ventricle wall.

**Fig.4.**
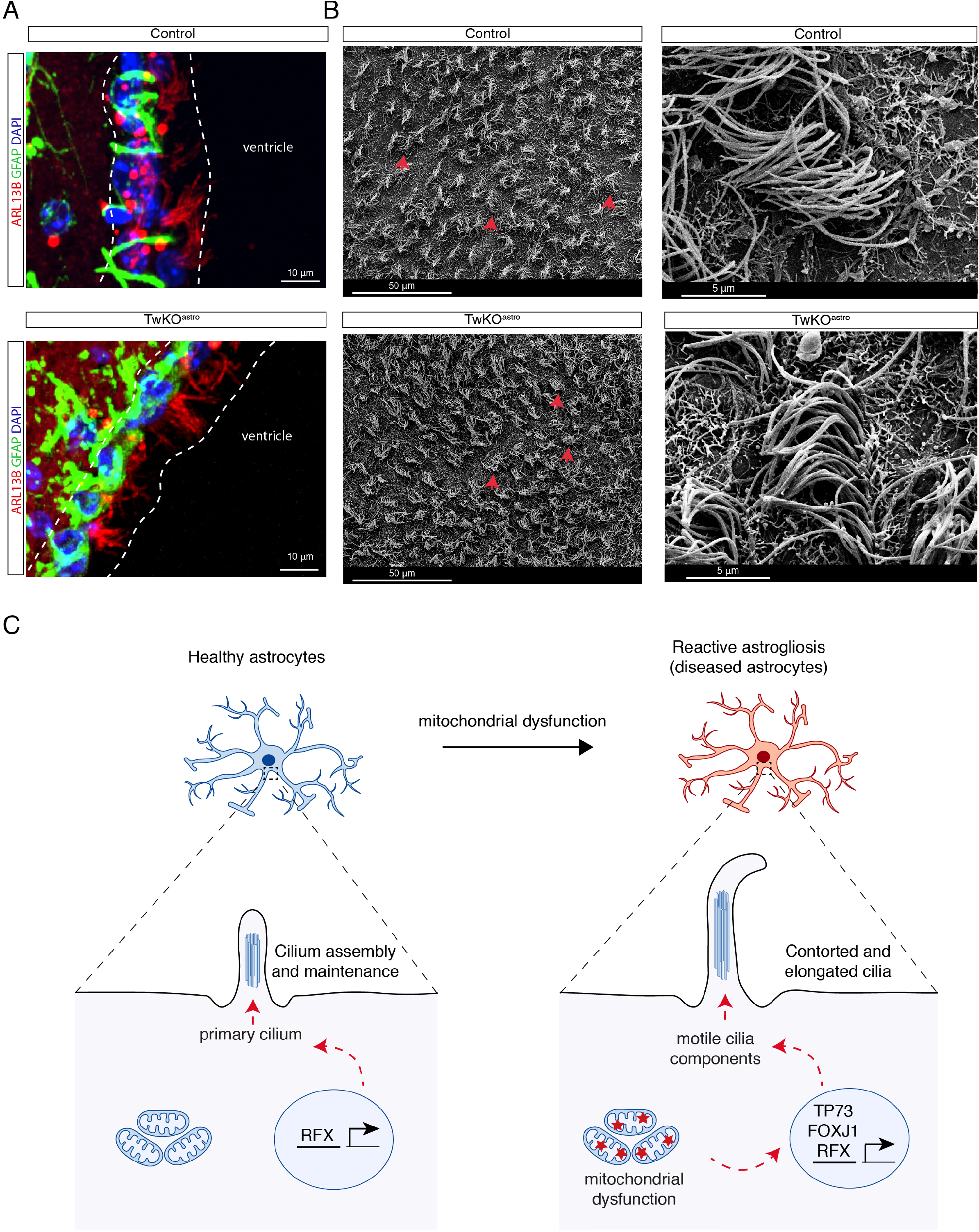
Ependymal cilia do not show apparent abnormalities upon mitochondrial dysfunction. (A-B): Cerebral cortex, 4.5-5 months old mice. (A) Ependymal cell layer (dashed lines) with astrocytes (GFAP^+^) and motile cilia (ARL13B^+^). (B) Tufts of cilia (red arrows) on the surface of ependymal cells. Scanning electron microscopy from ventricle side. (C) Mechanistic summary: Mitochondrial dysfunction induces a ciliogenic program in astrocytes. In a healthy setting, primary cilia are assembled during brain development upon cell cycle exit, and maintained in mature astrocytes. RFX factors regulate assembly of both primary and motile cilia, while FOXJ1 and TP73 are required for ependymal cell differentiation and formation of motile cilia. Mitochondrial dysfunction results in atypical Foxj1 induction in cortical astrocytes, coupled with expression of motile cilia components and contorted cilia morphology.

### Mitochondrial dysfunction in astrocytes alters brain lipid metabolism

Gene ontology analysis of downregulated transcripts in TwKO^astro^ sorted astrocytes indicated major changes in lipid metabolic pathways (four out of the top ten downregulated pathways) including cholesterol biosynthesis (Fig. S4A). Analysis of lipids and lipid-like molecules in an untargeted metabolomics dataset (Ignatenko et al., 2020) from TwKO^astro^ cortical lysates showed two ceramides and a cholesteryl ester depleted at the early 2.3-month-old timepoint, while in 3.2-month-old TwKO^astro^ mice 106 out of 172 identified lipid metabolites were changed (Fig. S4B, Suppl. Table 4). Specifically, the majority of identified steroids and derivatives, fatty acids and other fatty acyls, as well as prenol lipids were decreased; and sphingolipids, glycerolipids, and glycerophospholipids were imbalanced in amounts (Fig. S4B). Prominent changes were reflected also in lipid storage: ultrastructural analysis and biochemical detection with BODIPY^493/503^ probe revealed an accumulation of lipid droplets in TwKO^astro^ mice at late disease stage (five to eight months of age) (Fig. S4C, D). Recent reports propose that astrocytes import and metabolise lipids generated by neurons via mitochondrial fatty acid oxidation (Ioannou et al., 2019; Liu et al., 2015, 2017). A likely cause of accumulation of lipids in astrocytes is their impaired beta-oxidation as a consequence of mtDNA loss and oxidative phosphorylation deficiency. The lipids may, however, serve a function, as lipid droplets have been reported to be protective against lipotoxicity (Liu et al., 2015; Nguyen et al., 2017). Collectively, our data demonstrate that mitochondrial dysfunction in astrocytes induces a severe, progressive defect in brain lipid metabolism. Whether these changes affect the membrane protruding cilia and their homeostasis remains to be explored.

In conclusion, we show here that primary mitochondrial dysfunction in astrocytes induces structural aberration of primary cilia, the cellular antenna-like protrusions and hubs for nutrient and growth signaling. Our evidence indicates that these respiratory chain deficient astrocytes induce an anomalous expression of a motile cilia program (Figure 4), governed by transcription factors FOXJ1, and the RFX family. Despite the multiciliary program induction, TwKO astrocytes had single cilia, but their structure was elongated and sometimes extremely contorted. As underlying, potentially overlapping, mechanisms we propose 1) anomalously and chronically induced motile cilia factors trafficked to the primary cilium modifying the structure of the axoneme; 2) impaired turnover of ciliary proteins, as a consequence of ISR^mt^, a chronic anabolic stress response to mitochondrial dysfunction; 3) changed membrane structure because of major changes in cellular lipids.

The signaling axis between mitochondria and cilia has been recently suggested in studies of cultured cells, after toxin-mediated or genetic mitochondrial dysfunction (Bae et al., 2019; Burkhalter et al., 2019; Failler et al., 2020). Our in vivo evidence indicates that metabolic stress from mitochondrial respiratory chain dysfunction leads to a genetic rewiring of the ciliary pathway in differentiated cells. The potential signals include components of ISR^mt^, involving changes to levels of many critical metabolites such as amino acids, TCA cycle intermediates and nucleotides (Nikkanen et al., 2016), and anabolic metabolism related to ISR^mt^. Furthermore, here we show a major dysregulation of the astrocyte lipid profile as a result of defective mitochondrial fatty acid oxidation. If and how these candidate metabolites and lipids signal to induce a ciliogenic response will no doubt be the focus of future studies. These findings open completely new avenues for investigating metabolic ciliopathy as a contributor to pathophysiology of primary mitochondrial brain diseases of children and have potential relevance also for neurodegenerative pathologies associated with secondary mitochondrial dysfunction.

## Figure legends

**Fig. S1.**
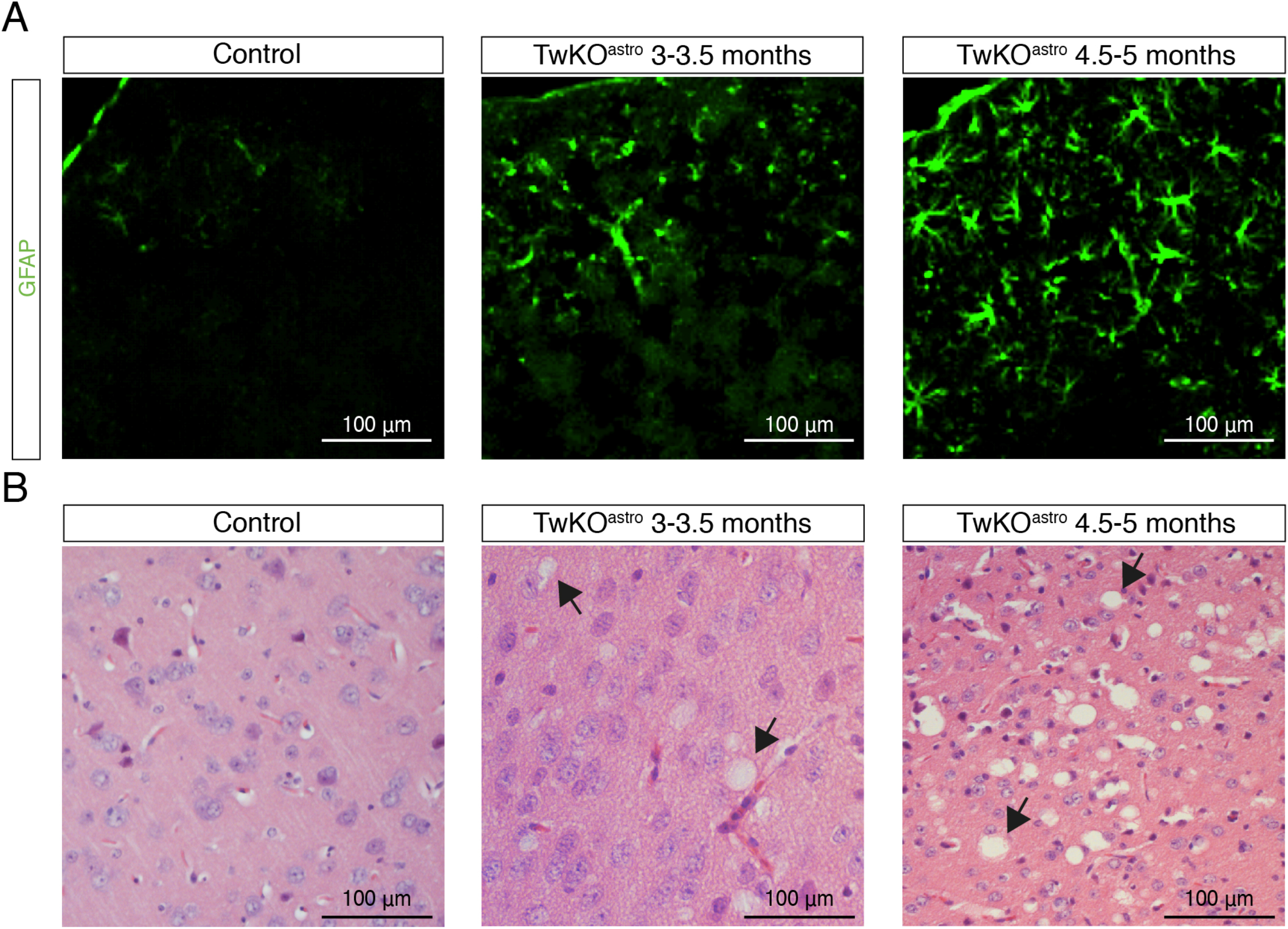
Related to Fig. 1 and 3. Histological consequences of Twinkle KO in mouse astrocytes. (A) Reactive gliosis in cerebral cortex of TwKO^astro^ mice. Immunostaining against GFAP. (B) Spongiotic encephalopathy in cerebral cortex TwKO^astro^ mice. Haematoxylin and eosin staining. Arrows indicate examples of spongiotic pathology.

**Fig. S2.**
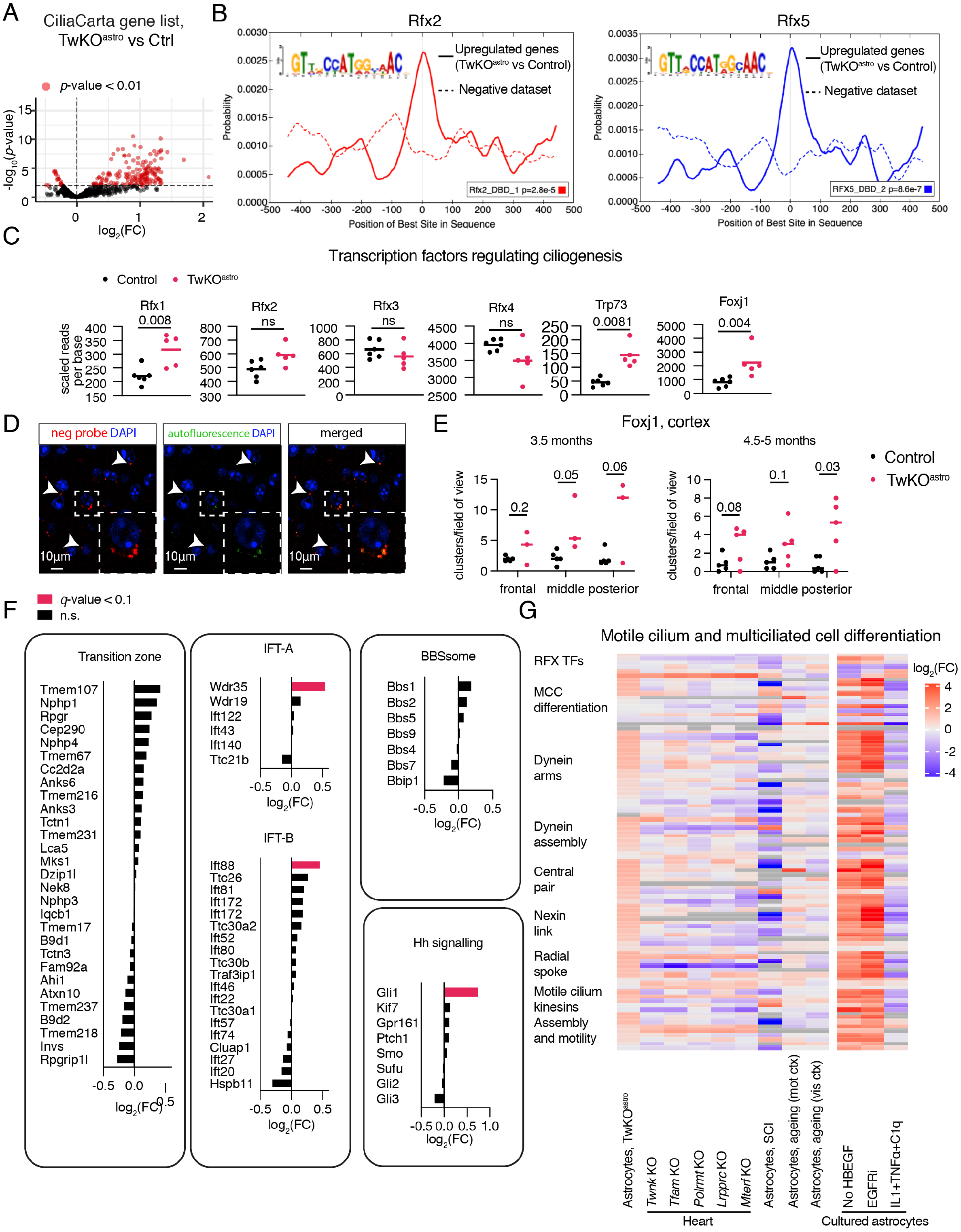
Related to Fig. 2. Transcriptomic consequences of Twinkle KO in isolated cortical astrocytes. (A) Expression levels of ciliary genes (CiliaCarta gene list (van Dam et al., 2019)). See also Fig. 2A. (B) Motif enrichment analysis of promoters of upregulated genes. Motif probability graph. See also Fig. 2C. (C) Expression of transcription factors that regulate ciliogenesis, scaled reads per base. Expression data for these genes are also presented in Fig. 2E-F. p-values calculated using unpaired two-tailed parametric t-test. (D) Control experiment for FoxJ1, RNA-fluorescence in situ hybridization with the negative probe. Arrowheads: puncta in the red channel (negative probe) that overlap with the green channel (autofluorescence) were excluded from analysis. Inset: an enlarged image. (Unrelated to this analysis, these mice also express AAV-gfaABC1D-Arl13b-eGFP). See also Fig. 2H. (E) Quantification of Foxj1 RNA-fluorescence in situ hybridization in the mouse brain cortex. Signal quantified from three fields of view per mouse, symbols represent the average per mouse. Clusters with a minimum of three puncta were quantified. Control n=5, TwKO^astro^ n=3. See also Fig. 2H. (F) Expression of pan-ciliary and specific to primary cilia factors in astrocytes sorted from TwKO^astro^ compared to control mice. See also Fig. 2J. IFT = intraflagellar transport; BBBsome = a component of the basal body; Hh signalling = hedgehog signalling pathway. (G) Regulation of motile ciliogenic program in astrocytes upon various insults and upon tissuespecific mitochondrial dysfunction. Datasets are from this study (described in Fig.1) and from (Anderson et al., 2016; Boisvert et al., 2018; Guttenplan et al., 2020; Kühl et al., 2017; Li et al., 2019). MCC = multiciliated cell; SCI = spinal cord injury; mot ctx = motor cortex; vis ctx = visual cortex. Expression data for some of these genes are also presented in Fig. E, J. (A-C, F): RNA sequencing, astrocytes purified from TwKO^astro^ compared to control mice. The dataset is described in Fig. 1. FC = fold change (A, F, G): See also Suppl. Table 2.

**Fig. S3.**
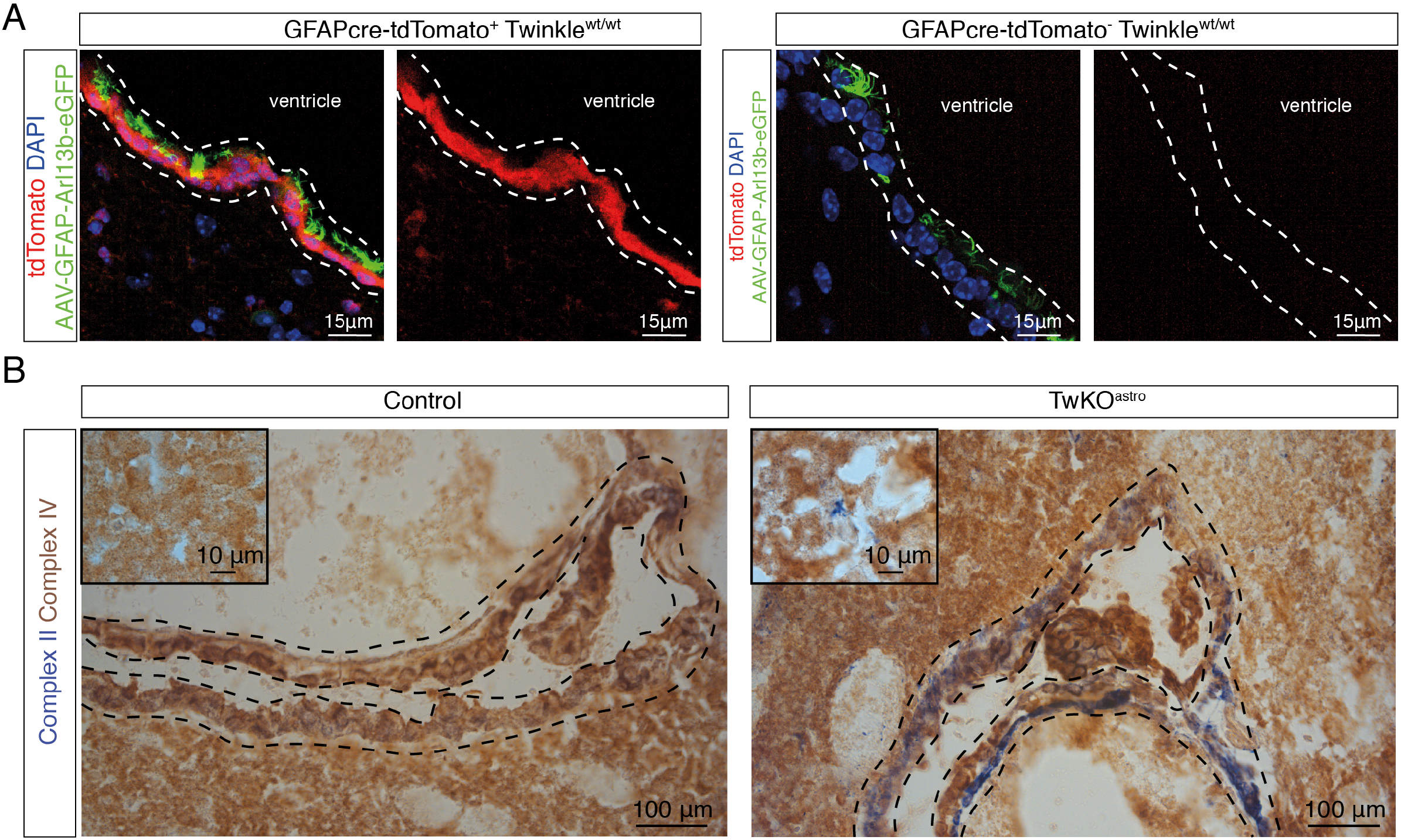
Related to Fig. 4. Cre-driver expression pattern and respiratory chain deficiency in TwKO^astro^ ependymal lining. (A) GFAP73.12-Cre expression, ventricular lining, mouse brain. Expression of tdTomato is controlled by a floxed stop cassette. Dashed lines indicate the ependymal cell layer. The mice also express AAV-gfaABC1D-Arl13b-eGFP to visualize cilia. (B) Histochemical in situ enzymatic activity assay of Complex IV (brown precipitate) and Complex II (blue) of OXPHOS; frozen brain sections, 5.5 months old mice. Ventricle, dashed lines, indicate ependymal cell layer. Insets are from the cortex. Blue areas represent increased Complex II (nuclear-encoded) and reduced Complex IV (mtDNA-encoded) activity, indicative of deficient mitochondrial OXPHOS function.

**Figure S4.**
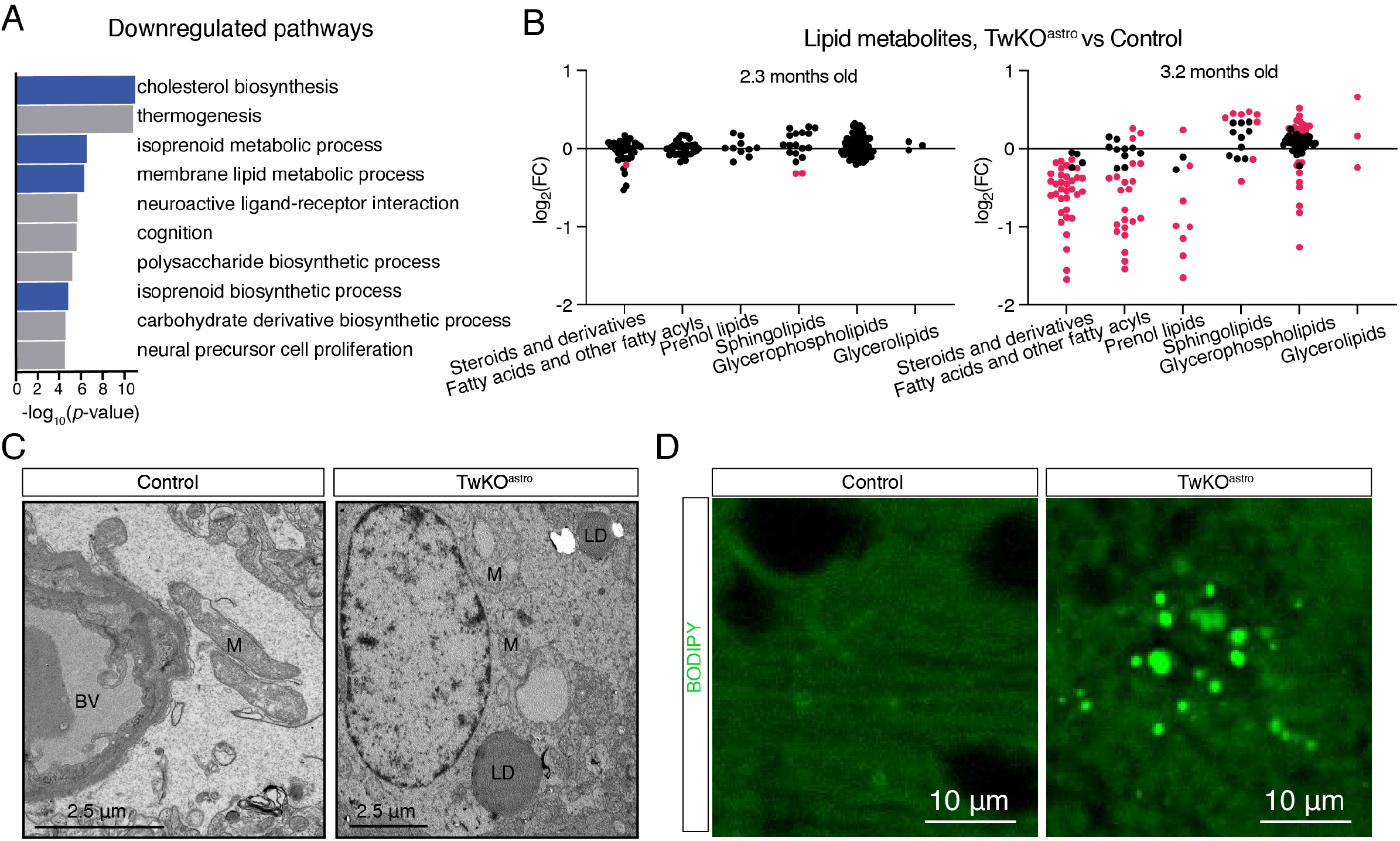
Related to Figure 1. Mitochondrial dysfunction in astrocytes alters brain lipid metabolism. (A) Gene ontology pathway enrichment analysis of downregulated genes. Blue colour denotes pathways related to lipid metabolism. (B) Lipids and lipid-like molecules, metabolomics (dataset from (Ignatenko et al., 2020)). See also Suppl. Table 4. FC = fold change. (C) Transmission electron microscopy, mouse brain cortex, 5-8 months old mice. LD = lipid droplet, M = mitochondria. (D) BODIPY^493/503^ neutral lipid dye staining, 5 months old mice, cortex.

## Supplementary tables

Suppl. Table 1. Reactive astrogliosis markers.

Suppl. Table 2. Cilia genes.

Suppl. Table 3. ChIP Seq analysis.

Suppl. Table 4. Lipids and lipid-like molecules.

Suppl. Table 5. Oligonucleotides used in the study.

Suppl. Table 6. RNA sequencing TwKO^astro^ vs Control purified adult astrocytes.

## Author contribution

O.I., G.I.D., and A.S. conceived and conceptualized the research. O.I. designed, planned and executed all main experiments, analysed and interpreted data. S.M. executed experiments, analysed and interpreted data. O.I., A.K., J.N., G.I.D executed and interpreted bioinformatic analyses. A.K. provided expertise for statistical analyses. H.V., E.J supervised electron microscopy experiments. O.I. assembled figures and wrote the first draft of the manuscript. O.I., G.I.D. and A.S. wrote the manuscript, and all authors commented. G.I.D. and A.S. supervised the study.

## Acknowledgements

We thank Brendan Battersby and Maxim Bespalov for insightful discussions; Kirsi Mattinen, Markus Innilä, Mervi Lindman, Tuula Manninen, Babette Hollmann, Sonja Jansson, and Markus Innilä for technical assistance; Cory Dunn for Arl13b-eGFP plasmid. We acknowledge the following core facilities of the University of Helsinki for their resources and expertise: Biomedicum Imaging Unit, Genome Biology Unit, AAV core, and Laboratory Animal Center.

We thank the following funding sources: Academy of Finland, Sigrid Juselius Foundation, Jane and Aatos Erkko Foundation, University of Helsinki (A Suomalainen); University of Helsinki Doctoral Program In Biomedicine, Biomedicum Foundation, Otto Malm Foundation (O Ignatenko); European Molecular Biology Organization (ALTF 1185-2017), Human Frontier Science Program Organization (LT000446/2018-L) (J Nikkanen). The authors declare no competing financial interests.

## Materials and Methods

### Animal experimentation

Animal experiments were approved by The National Animal Experiment Review Board and Regional State Administrative Agency for Southern Finland, following the European Union Directive. Mice were maintained in a vivarium with 12-h light:dark cycle at 22 °C and allowed access to food and water ad libitum. TwKO^astro^ mice (Ignatenko et al., 2018) (Gfap73.12Cre^+^; Twnk^loxp/loxp^ mice) were generated by breeding mice carrying floxed Twnk alleles (Twnk^loxp/loxp^ or Twnk^+/loxp^) to Gfap73.12-Cre^+^ mice (JAX: 012886) creating a deletion of exons 2 and 3 of Twnk. Littermates were used as controls. Twnk^loxp/+^, Twnk^loxp/loxp^, Gfap73.12Cre^+^; Twnk^loxp/+^, Gfap73.12Cre^+^; Twnk^+/+^ were used as controls. The methodology for generating Twnk^loxp/loxp^ mice is previously described (Nikkanen et al., 2016). These mice carried Y508C mutation in the targeted Twnk gene (Nikkanen et al., 2016). Using the same method as in (Nikkanen et al., 2016), for this study we regenerated Twnk^loxp/loxp^ mice without the Y508C mutation. Gfap73.12Cre^+^;Twnk^loxp/loxp^ and Gfap73.12-Cre^+^;Twnk^Y508C/Y508C^ mice were indistinguishable in mtDNA brain pathology and were used interchangeably as TwKO^astro^. Cre reporter mice were generated using Ai14 tdTomato mice (JAX: 007914). Mice on C57Bl/6OlaHsd or mixed genetic background mice were used. For all experiments mice were either terminally anesthetized by intraperitoneal injection of pentobarbital or were euthanized with CO_2_.

### Astrocyte sorting from adult mice

Astrocytes were purified using magnetic beads coated with the ACSA-2 antibody, similar to manufacturer’s instructions and as previously described (Batiuk et al., 2017; Holt et al., 2019). Cerebral cortex samples from four mice per sorting preparation were dissected. Tissue was dissociated using an enzymatic kit (Adult Brain Dissociation (P) Kit, Miltenyi Biotec, #130-107-677). Myelin debris were depleted using Myelin Removal Beads II (Miltenyi Biotec, #130-096-731). Astrocyte fraction was enriched using Anti-ACSA-2 MicroBead Kit (Miltenyi Biotec, #130-097-679).

### Brain collection for histological analyses

Mice were transcardially perfused with ice-cold PBS followed by perfusion with ice-cold 4 % formaldehyde solution in PBS. The brains were postfixed in 4 % formaldehyde solution in PBS overnight at 4 °C. For immunofluorescence, brains were stored in PBS with 0.02 % sodium azide at 4 °C and then for several days incubated in 30 % sucrose in PBS solution before freezing. For RNA-fluorescence in situ hybridisation, after postfixing brains were transferred to 30 % sucrose in PBS for several days at 4 °C, and then frozen. Freezing was done in embedding media (O.C.T. compound Tissue-Tek #4583) at −20 °C. The brain sections were made using Thermo Scientific Cryostar NX70 cryostat at 12-20 μm to adhesive microscope slides (Matsunami Glass #TOM-11) or ThermoFisher Scientific (J1800BMNZ).

### Immunofluorescence

Mouse brain sections or coverslips with cultured astrocytes were incubated in 10 % horse serum, 0.1 % Triton X-100 (or 1.0 % Tween20) in PBS for one hour at room temperature. Samples were then incubated overnight at +4 °C with primary antibodies diluted in the same solution or in an antibody diluent (Agilent #S3022). The samples were then washed in 0.1 % Triton X-100 or 1.0 % Tween20 in PBS for 30-60 minutes. Samples were then incubated for one hour at room temperature with secondary antibodies conjugated with Alexa Fluor fluorescent probes (Thermo Fisher Scientific) diluted in 10 % horse serum, 0.1 % Triton X-100 or 1.0 % Tween20 in PBS and mounted with DAPI-containing mounting medium (Vectashield #H-1200-10). Images were acquired using the Andor Dragonfly spinning disk confocal microscope. Maximum intensity projection images are presented in figure panels. Images were also acquired with a Zeiss Axio Imager epifluorescent microscope. For all images, only linear adjustments were applied. Antibodies used in the study: anti-GFAP (Sigma-Aldrich AB5804, 1:500 for mouse brain sections, 1:1000 for cultured cells); anti-Arl13b (Abcam #AB136648, 1:500 for mouse brain sections, 1:1000 for cultured cells; anti-Arl13b (Proteintech #17711 −1-AP, 1:1000 for cultured cells). Maximum intensity projections of confocal images are shown.

### RNA-fluorescence *in situ* hybridisation

Frozen mouse brain sections were dehydrated with ethanol, and hybridised with a probe against mouse Foxj1 according to the manufacturer’s instructions (ACD Systems #317091, #323110 and PerkinElmer #FP1487001KT, #FP1488001KT). Images were acquired using Andor Dragonfly spinning disk confocal microscope, 4 μm stacks with a 1 μm step were acquired. Signal was manually quantified from maximum intensity projection images using Fiji software by an investigator blind to genotypes. Clusters of a minimum of three puncta were quantified.

The signal was quantified from three fields of view per mouse. Maximum intensity projections of confocal images are shown.

### Lipid staining

To visualise lipid droplets on mouse brain sections, BODIPY^493/503^ dye (intrinsic affinity to neutral and non-polar lipids) was used according to manufacturer’s instructions (Thermo Fisher Scientific #D3922). Mice were transcardially perfused with ice-cold PBS, followed by perfusion with ice-cold 4 % formaldehyde solution. Brains were postfixed overnight, incubated for several days in 30 % sucrose and frozen in the embedding media and sectioned. Sections were incubated for 10 minutes in PBS, 10-60 minutes in BODIPY solution (1:100-1:1000 of the stock solution in PBS), washed twice for five minutes in PBS, mounted, and imaged.

### Analysis of cilia morphology

Cilia morphology was analysed on images acquired with Andor Dragonfly spinning disk confocal microscope (7 μm stacks with a 0.5 μm step) using Imaris 9.5.1 software with Surfaces module. Settings for surfaces were adjusted to automatically detect cilia, which was followed by manual selection of surfaces of interest. Arl13b^+^ signal in Aldh1l1^+^ cells was analysed. Object oriented bounding box length C length was used as a proxy of ciliary length. Categories of morphology of cilia were assigned manually, using three-dimensional images. Cilia were analysed from six fields of view per mouse.

### Transmission electron microscopy

Mice were euthanised using CO_2_. 1-2 mm^3^ pieces of cortex were dissected out and fixed overnight at 4 °C with 2.5 % glutaraldehyde (EM-grade, Sigma-Aldrich check in the lab) in 0.1 M sodium phosphate buffer, pH 7.4. After washing, samples were post-fixed with 1% non-reduced osmium tetroxide in 0.1 M sodium phosphate buffer for 1 h at room temperature, dehydrated in ethanol series and acetone prior to gradual infiltration into Epon (TAAB 812, Berks, UK). After polymerization at 60 °C for 18 h, ultrathin sections were cut and post-stained with uranyl acetate and lead citrate. TEM micrographs were acquired with a Jeol JEM-1400 microscope (Jeol Ltd, Tokyo, Japan) running at 80 kV using an Orius SC 1000B camera (AMETEK Gatan Inc., Pleasanton, CA, USA).

### Scanning electron microscopy

Mice were transcardially perfused with ice-cold PBS followed by perfusion with ice-cold 4 % formaldehyde solution in PBS. Brains were postfixed in 2 % formaldehyde solution in 0.1 M sodium phosphate buffer for 2-6 more hours at room temperature. En-face whole-mounts of the lateral ventricle wall were dissected as previously described (Mirzadeh et al., 2010), and postfixed in 2.5 % glutaraldehyde in 0.1 M sodium phosphate buffer. Samples were stained in 2 % non-reduced osmium tetroxide in 0.1 M sodium phosphate buffer, pH 7.4 for 2 hours, dehydrated in ascending concentrations of ethanol, and dried by critical point drying (Leica EM CPD300, Leica Mikrosystems, Wetzlar, Germany). The samples were oriented using dissection microscope, mounted on aluminum stubs covered with carbon tape, and coated with a thin layer of platinum. Scanning electron microscopy micrographs were taken under high vacuum using a FEI FEG-SEM Quanta 250 (Thermo Fisher Scientific, Waltham, MA, USA) with a 5.00 kV beam, spot size 3.5.

### *In vivo* labelling of cilia

To label cilia in vivo in the mouse brain, pups (P0-P5) were injected intraventricularly with AAV-gfaABC1D-Arl13b-eGFP viral vector. pAAV-ABC1D-Arl13b-eGFP plasmid was subcloned using destination vector pAAV-GFAP-GFP (addgene 50473), and a PCR-amplified insert with overhangs complimentary to destination vector from pLi3-Arl13b-eGFP plasmid (addgene 40879 (Larkins et al., 2011)). Assembly was done using NEB HiFI DNA assembly kit (NEB E5520) according to manufacturer’s instructions. The plasmid was validated by direct sequencing of the insert fragment. For AAV production, the plasmid was amplified using chemically competent cells deficient for recombinase activity (Agilent Technologies #200152), purified (Macherey-Nagel #740416.10), and packaged into AAV8 vector (packaging was done by University of Helsinki AAV core unit). For injections, pups were cryoanesthetized and injections were done using stereotaxic frame as previously described (Kim et al., 2014); 2 ul per hemisphere was administered, the AAV titre was 3-3.5×10e-9 vp/μl.

### In situ enzyme activity of OXPHOS Complexes

The histochemical activity assay was done as described previously (Forsström et al., 2019). Mice were euthanised with CO_2_, freshly collected brains were embedded in embedding media (O.C.T. compound Tissue-Tek #4583) and frozen in 2-methylbutane bath under liquid nitrogen cooling. For simultaneous colorimetric detection of the activity of Complex IV (cytochrome-c-oxidase (COX)) and Complex II (succinate dehydrogenase (SDH)), 12 μm brain sections were incubated with enzyme substrates (30’ for COX, room temperature), 40 minutes for SDH (+37 °C). COX: 0.05 M phosphate buffer (pH 7.4) with 3,3–diaminobenzidine (DAB), catalase, cytochrome c and sucrose. SDH: 0.05 M phosphate buffer (pH 7.4) with nitro-blue tetrazolium and sodium succinate. Sections were then dehydrated by incubations in ascending concentrations of ethanol, xylene-treated and mounted.

### RT-qPCR

RNA from purified astrocytes or brain cell suspension after enzymatic digestion was extracted using RNA-binding columns (NucleoSpin RNA plus Macherey-Nagel #740984.250), genomic DNA was eliminated and cDNA was synthetised (Thermo Fisher Scientific #K1672). Real-time quantitative PCR (RT-qPCR) reactions were performed (Bioline #BIO-98020). NCBI primer BLAST software was used for oligonucleotide design (Ye et al., 2012), 70–150 bp-long product was amplified, and oligonucleotides were designed to be separated by at least one intron of the corresponding genomic DNA of a minimum 1,000 bp length (whenever possible). Oligonucleotides for amplification of Atp1b2, Slc1a3, Cspg4, Ocln are from (Batiuk et al., 2017) and Syt1 from (Liddelow et al., 2017). Oligonucleotide sequences can be found in Suppl. Table 5. The expression of the genes of interest was normalized to hydroxymethylbilane synthase (Hmbs or Ywhaz) expression.

### mtDNA quantification

Purified astrocytes or brain tissue suspension after enzymatic digestion was lysed in TNES buffer with Proteinase K overnight at 55 °C water bath. DNA was extracted using phenol-chloroform, followed by ethanol and ammonium acetate precipitation. Pellet was washed with ethanol, dried, and resuspended in 10 mM Tris-Cl, pH 8.0. Quantitative analysis of mtDNA was performed by quantitative PCR, normalized to a nuclear gene Rmb15 as described in (Ignatenko et al., 2020). Oligonucleotide sequences can be found in Suppl. Table 5.

### Metabolomics

Metabolomics datasets are from (Ignatenko et al., 2020). All metabolites belonging to lipids and lipid-like molecules (HMDB super class) were analysed and grouped for plotting according to HMDB classification. The mice were 2.3 months old. See also Suppl. Table 4.

### RNA sequencing

RNA was extracted from purified astrocyte preparations using RNA-binding columns (NucleoSpin RNA plus Macherey-Nagel #740984.250). RNA was analyzed using TapeStation, and only samples with RIN>7 were used. RNA sequencing was done in the BGI Genomics, China. mRNA was enriched by Oligo dT selection, and reverse transcription was done using random N6 primer. End repair and adaptor ligation were followed by PCR amplification of the library. The library was sequenced using BGISEQ-500 platform, paired-end sequencing, read length 100 bp. After sequencing, reads were filtered to remove adaptor sequences, contamination, and low-quality reads using Soapnuke software version 1.6.7 version (filtering parameters -n 0.001 -l 20 -q 0.4 -A 0.25). Additionally, quality of reads was analysed using FastaQC software. The data are available in NCBI GEO repository (accession number GSE174343) (Edgar et al., 2002). See also Suppl. Table 6.

### Bioinformatic analyses

Quantification of transcript abundance in RNA sequencing dataset was done using Kallisto software (mm10 build, (Bray et al., 2016)). Differential expression analysis and PCA plot were prepared with Sleuth (Pimentel et al., 2017), gene ontology pathway enrichment analysis was done with with Metascape (Zhou et al., 2019). Upregulated genes: top 1000 gene were analysed (genes with q-value<0.1 and log_2_(FC)>0.3 were selected, and then sorted by log_2_(FC)). Downregulated genes: all genes with q-value<0.1 and log_2_(FC)<−0.3 were selected (408 genes). RNA sequencing analysis of published datasets (Suppl. Table 2): available from NCBI GEO repository (GSE76097; GSE99791; GSE96518; GSE143598; GSE125610). log_2_(FC) were used (Anderson et al., 2016; Boisvert et al., 2018; Guttenplan et al., 2020; Kühl et al., 2017), or log_2_FC were calculated from raw read counts using DeSeq2 (Li et al., 2019; Love et al., 2014).

For motif enrichment analysis (Suppl. Table 3), genomic sequences corresponding to 500 bp upstream and 500 bp downstream from the transcription start sites (TSS) of upregulated genes were extracted (log_2_(FC)>0.25 and p-adj-value<0.05) (mm10 build) (Yates et al., 2020). As a negative dataset, 1,000 randomly selected genes from the mouse genome were used. To find enriched motifs CentriMo tool from the MEME Suite was used (Bailey and Machanick, 2012). For analysis of TF binding site enrichment, genomic coordinates for binding sites of RFX1 −3 based on a previously published ChIP-Seq study were used (Lemeille et al., 2020). Genomic coordinates spanning 10,000 bp upstream and 10,000 bp downstream from TSSs of upregulated genes (log_2_(FC)>0.25 and p-adj-value<0.01) were extracted. An overlay of the two datasets in Excel revealed the RFX binding sites within TSS proximal sequences of upregulated genes. As negative datasets either randomly selected genes from the mouse genome or genes which were expressed in astrocytes but not changed based on our RNA-Seq results were used. GRCm38 (mm10) mouse genome build was used for all analyses and manipulations on genomic intervals were carried out using the Galaxy platform (Afgan et al., 2018).

### Statistical analysis

Statistical analyses were performed using Prism 8 software and R; graphs were made with Prism 8. Statistical analysis of differential gene expression of RNA sequencing data generated in this study was done using the Sleuth package (Pimentel et al., 2017), q-values were used throughout the study to determine statistical significance of these comparisons. Statistical analyses of the cilia length distribution were done using Kolmogorov-Smirnov test in Prism 8 and ANOVA test in R base software. Statistical analysis of cilia morphology and RNA-fluorescence in situ hybridization between two groups was performed using unpaired twotailed parametric t test in Prism 8. Heatmaps were generated using Complex Heatmap package in R (Gu et al., 2016), Z-scores were calculated from scaled reads per base.

